# OptDesign: Identifying Optimum Design Strategies in Strain Engineering for Biochemical Production

**DOI:** 10.1101/2021.12.10.472123

**Authors:** Shouyong Jiang, Irene Otero-Muras, Julio R. Banga, Yong Wang, Marcus Kaiser, Natalio Krasnogor

## Abstract

Computational tools have been widely adopted for strain optimisation in metabolic engineering, contributing to numerous success stories of producing industrially relevant biochemicals. However, most of these tools focus on single metabolic intervention strategies (either gene/reaction knockout or amplification alone) and rely on hypothetical optimality principles (e.g., maximisation of growth) and precise gene expression (e.g., fold changes) for phenotype prediction. This paper introduces OptDesign, a new two-step strain design strategy. In the first step, OptDesign selects regulation candidates that have a noticeable flux difference between the wild type and production strains. In the second step, it computes optimal design strategies with limited manipulations (combining regulation and knockout) leading to high biochemical production. The usefulness and capabilities of OptDesign are demonstrated for the production of three biochemicals in *E. coli* using the latest genome-scale metabolic model iML1515, showing highly consistent results with previous studies while suggesting new manipulations to boost strain performance. Source code is available at https://github.com/chang88ye/OptDesign.

## Introduction

A growing population and fast economical development are leading to an increasing demand of various daily products and industrial raw materials, many of which are derivatives of oil and petroleum. Over the past decades, important efforts are being made to develop sustainable production processes that convert biomass or other renewable resources to bio-products through cell platforms. ^1^ A key challenge in this respect is the design of high-performance strains with efficient metabolic conversion routes to desired products. Recent advances in genome-scale metabolic modelling (GSMM)^2^ has made it possible to have a system-level understanding of cell physiology and metabolism, leading to rational prediction of metabolic interventions for strain development. Systems strain design^1^ has helped to improve the production of numerous biochemicals, including lycopene, ^3^ malonyl-CoA, ^4^ alkane and alcohol,^5^ and hyaluronic acid.^6^

A number of tools have been developed for strain design.^7,8^ OptKnock,^9^ which was developed to block some reactions in metabolic networks, is one of the earliest such tools. OptKnock identifies the knockout targets that leads to maximal biochemical production in the context of flux balance analysis^2^ that is subject to mass balance and thermodynamic constraints. This results in a bilevel optimisation problem which can be solved through mathematical reformulation into a standard mixed-integer linear program (MILP). ^9^ The OptKnock model was latter extended to consider gene up/down-regulation,^10^ swap of cofactor specificity,^11^ and introduction of heterologous pathways^12^ for biochemical production. It was also adapted to identify synthetic lethal genes for anti-cancer drug development.^13^ Some improvement strategies, such as GDBB^14^ and GDLS,^15^ have been proposed to improve the efficiency of OptKnock in solving the bilevel problem. There also exist numerous approximate solutions to the OptKnock model, including genetic algorithms^16^ and swarm intelligence.^17^ Designing strains that couple production to growth has received increasing attention in recent years, mainly due to the great production potential of growth-coupled strains in adaptive laboratory evolution.^18^ Consequently, a number of computational tools along this direction have been developed to design strains with various growth-coupled phenotypes.^19,20^ OptCouple^20^ simulates jointly gene knockouts, insertions, and medium modifications to identify growth-coupled designs, although gene expression regulation is not considered. In addition, game theory has been introduced into metabolic engineering.^21,22^ NIHBA^22^ considers metabolic engineering design as a network interdiction problem involving two competing players (host strain and metabolic engineer) in a max-min game enabling growth coupled production phenotypes, and the problem is solved by an efficient mixed-integer solver. Furthermore, there are also some studies which do not rely on optimality principles for phenotype prediction. Among these, the minimum cut set (MCS) based approach, ^23,24^ which aims to find the smallest number of interventions blocking undesired production phenotypes, has been extensively studied. Despite high computational complexity, MCS-based approaches have successfully predicted strain design strategies leading to *in vivo* biochemical production. ^25^ Another important approach of the same kind is OptForce, which identifies metabolic interventions by exploring the difference in flux distributions between the wild type and the desired production strain.^26^ OptForce has showed good predictions for *in vivo* malonyl-CoA production.^4^

The use of computational tools is undisputedly important to strain development in metabolic engineering.^27^ However, there are several limitations which may prevent the wide applicability of the above-mentioned approaches. First, most of the tools focus on prediction of either knockout targets or regulation targets alone, with a few exceptions that are capable of predicting both interventions, such as OptForce^26^ and OptRAM.^28^ These exceptions highlight that a combination of knockout and up/down-regulation often leads to higher biochemical production compared to a single strategy. OptForce encourages the use of flux measurements while identifying optimum design strategies. OptRAM considers regulatory networks from which transcriptional factors can be optimised for biochemical production. However, both OptForce and OptRAM rely heavily on precise expression level of regulation targets; for example, desired production phenotypes can only be achieved at the exactly suggested flux values (OptForce) or up/down-regulation fold changes (OptRAM). It is known that gene expression is a complex process with many uncertainties. The underlying strict expression requirements in these approaches may miss theoretically non-optimal but practically feasible design strategies. In addition, both approaches rely on a reference flux vector of the wild type, which can be incorrectly chosen from many steady-state flux distributions if it cannot be uniquely determined. Second, many existing strategies rely on the assumption of optimality principles, e.g. maximal growth in OptKnock^9^ and derivatives, in cell metabolism. However, this assumption is not always an accurate representation of how cells respond to metabolic perturbations or environmental changes.^29^ NIHBA^22^ showed that reducing uncessary surrogate biological objectives helps to identify many non-optimal but biologically meaningful knockout solutions. Also, as noted in OptForce,^26^ most existing methods do not pro-actively integrate flux measurements into strain design for better phenotype prediction. For example, tools like OptKnock implicitly assume that the wild-type and mutant strains have the same flux space. The mutant flux space may be incorrectly constrained if it is not a subspace of the wild-type flux measurements.

This paper introduces a new computational tool, called OptDesign, that uses a two-step strategy to predict rational strain design strategies for biochemical production. OptDesign has the following capabilities:

**(C1)** overcomes uncertainty problem as there is no assumption of exact fluxes or fold changes that cells should have for production. As a result, non-optimal but good feasible solutions are not missed.
**(C2)** allows two types of interventions (knockout and up/down regulation).
**(C3)** does not assume (potentially unrealistic) optimal growth in production mode.
**(C4)** pro-actively integrates flux measurements if available.
**(C5)** does not need reference flux vector.
**(C6)** considers growth-coupled production.

OptDesign is the only tool that combines these six capabilities, as shown in Table 1. In the remainder of this paper, we describe OptDesign and benchmark it considering three case studies, demonstrating high consistencies of predicted design strategies with previous *in vivo* and *in silico* studies.

**Table 1:** A comparison of different strain design tools.

| Tool | C1 | C2 | C3 | C4 | C5 | C6 |
| --- | --- | --- | --- | --- | --- | --- |
| OptKnock<sup>9</sup> | ✗ | ✗ | ✗ | ✓ | ✓ | ✗ |
| OptForce<sup>26</sup> | ✗ | ✓ | ✗ | ✓ | ✗ | ✗ |
| OptCouple<sup>20</sup> | ✗ | ✗ | ✗ | ✗ | ✓ | ✓ |
| OptReg<sup>10</sup> | ✗ | ✓ | ✗ | ✗ | ✓ | ✗ |
| OptRAM<sup>28</sup> | ✗ | ✓ | ✗ | ✓ | ✗ | ✗ |
| NIHBA<sup>22</sup> | ✓ | ✗ | ✓ | ✗ | ✓ | ✓ |
| OptDesign (this study) | ✓ | ✓ | ✓ | ✓ | ✓ | ✓ |

## Materials and Methods

A metabolic network of *m* metabolites and *n* reactions has a stoichiometric matrix *S* that is formed by stoichiometric coefficients of the reactions. Let *J* be a set of *n* reactions and *v*_*j*_ the reaction rate of *j* ∈ *J, Sv* represents the concentration change rates of the *m* metabolites. The flux space *FS* is defined as the space spanned by all possible flux distributions *v* for the system subject to thermodynamic constraints at steady state (i.e, the concentration change rate is zero for all the metabolites). Mathematically, *FS* can be described as:

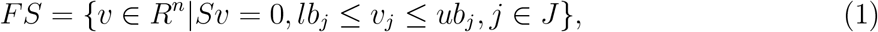

where *lb*_*j*_ and *ub*_*j*_ are the lower and upper flux bounds of reaction *j*, respectively. We use the notation *FS*_*w*_ for the wild type and *FS*_*m*_ for the mutant strain. Flux balance analysis (FBA) determines a single solution in *FS* when a surrogate biological objective is provided for (1).

OptDesign recognises metabolic changes from the wild type to production (mutant) strains. Let *v* ∈ *FS*_*w*_ denotes a flux vector of the wild type and Δ*v* denotes the flux change needed for *v* to transition into a desired production state. Obviously, *v* + Δ*v* represents a flux vector of the production strain, and it needs to satisfy mass balance and some production requirements, i.e., *v* + Δ*v* ∈ *FS*_*m*_. Note that flux measurements can be used to customise the flux bounds in *FS*_*w*_ and *FS*_*m*_ if available; otherwise, the flux bounds can be set according to flux variability analysis (FVA) predictions. For example, *FS*_*m*_ can be constrained by imposing production requirements on the lower bounds of the production reaction and biomass.

OptDesign introduces the concept of noticeable flux difference *δ* between the wild type strain and the production strain in reactions. OptDesign uses this concept to identify an optimal set of manipulations leading to the production phenotype *FS*_*m*_. To do so, OptDesign performs two key steps of optimisation. First, OptDesign identifies a minimal set of reactions that must deviate from their wild-type flux with at least *δ* in order to achieve *FS*_*m*_. This set of reactions form candidate regulation targets. Second, OptDesign searches through regulation candidates, together with knockout candidates, for the optimal combination of manipulations to maximise biochemical production. The following two subsections are devoted to presenting these two steps in detail.

### Selecting up/down-regulation reaction candidates

This step of OptDesign is to identify the minimum number of reactions whose flux must have a noticeable change if cellular metabolism shifts from the wild type to the required production state. A reaction is considered a candidate for up-regulation if its flux in the mutant is at least *δ* units more than that in the wild type. On the contrary, this reaction is considered for down-regulation if its flux in the mutant is at least *δ* units fewer than that in the wild type. Note that the above directional up/down-regulation definition is used for computational convenience, and final regulation targets identified from OptDesign will be rationally grouped by contrasting the wild type to the mutant strain by their absolute flux values (which will be detailed later in section Materials and Methods). In any other situations, this reaction is not considered as a candidate for genetic manipulation. Fig. 1 illustrates the above concept with a toy network of five reactions. Suppose *δ* is set to 2 units for all these five reactions, R4 and R5 are considered for down-regulation and up-regulation respectively. However, R1, R2 and R3 are not selected as regulation candidates since their flux changes from the wild type to the mutant are within the predefined threshold *δ*.

**Fig. 1.**
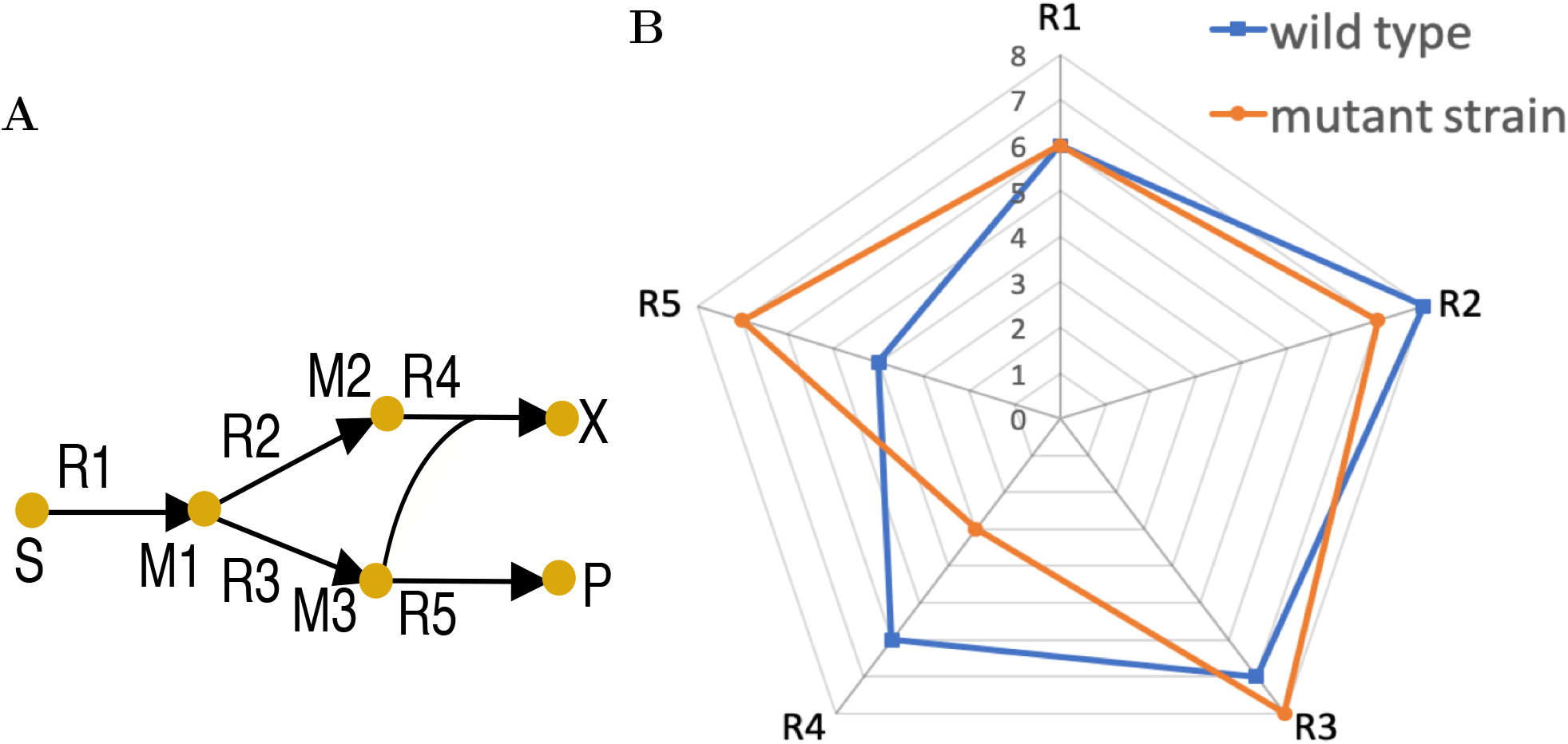
Toy metabolic network (A) and flux distributions of the wild type and mutant (B). Symbols in Fig. 1(A) are as follows: S, carbon source; X, biomass; P, product; M*i* (*i* = 1, 2, 3), metabolite name; R*i* (*i* = 1, …, 5), reaction name. Each axis in Fig. 1(B) represents the absolute flux for a reaction.

An MILP procedure is employed to minimise the number of reactions that must change their flux from the wild type to the mutant by at least *δ* units. This can be expressed as the following MILP problem:

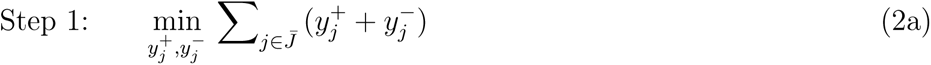

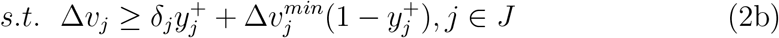

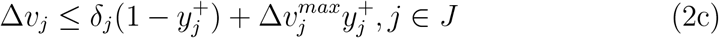

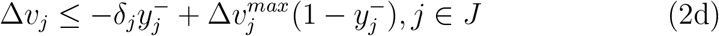

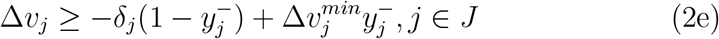

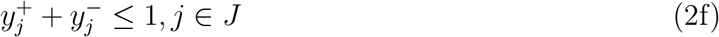

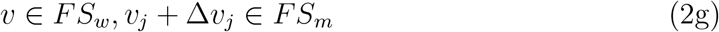

where 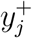 and 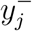 are binary variables representing the flux of reaction *j* increases and decreases by at least a noticeable level *δ*_*j*_ *>* 0 from the wild type to the production phenotype, respectively. Equivalently, 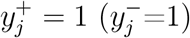 implies Δ*v*_*j*_ ≥ *δ*_*j*_ (Δ*v*_*j*_ ≤ −*δ*_*j*_). Constraints (2b)-(2c) and (2d)-(2e) are for flux increase and decrease, respectively. 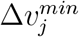 and 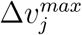 are the lower and upper bounds of flux change Δ*v*_*j*_, respectively. Special reactions in which fluxes are not allowed to be decreased (increased), e.g., non-growth associated maintenance, should have a zero value for the lower (upper) bound of their corresponding Δ*v* components. Each reaction cannot increase and decrease flux simultaneously, which implies the constraint (2f). Constraints (2g) describe the flux space of the wild type and mutant strain at steady state.

Note that this procedure does not need a known reference flux vector for both the wild type and the mutant, instead it takes into account all possible wild-type and mutant flux distributions that meet engineering requirements (e.g., growth rate, production/yield). However, it is recommended to make use of flux measurements for the wild-type and mutant strains if possible, in order to select a rational set of regulation targets effectively.

### Identifying optimal manipulation strategies

The solution to the MILP (2) results in a flux-increase set *F* ^+^ (corresponding to reactions with 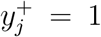) and a flux-decrease set *F* ^−^ (corresponding to reactions with 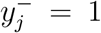), in addition to the suggested flux change Δ*v*. However, these two sets are not the minimum number of manipulations needed for the required production state as the effects of some manipulations can be propagated to the whole metabolic network.^26^ In addition, there is an engineering cost in manipulating reactions (through gene-protein-reaction associations), and therefore it is assumed there is a limit on the number *K*_*m*_ of genetic manipulations, including up/down-regulation and knockout. We allow gene/reaction knockout in this step for two reasons. Firstly, reactions having an unnoticeable flux change (below *δ*_*j*_ and thus not in *F* ^+^ or *F* ^−^) may sometimes be good manipulation targets, especially when there are involved in completing pathways. Taking them as potential knockout targets could improve biochemical production (essentially a relaxation of optimisation models). Secondly, there may be reactions in the regulation candidate set that carry near-zero fluxes in the production strain. From a practical point of view, completely deactivating them by gene knockout is easier than regulating their gene expression precisely to the suggested minute fluxes. Reaction knockout candidates (denoted by set *F* ^×^) can be selected by a preprocessing approach,^22^ which excludes reactions that are essential, irrelevant or unlikely to be good knockout targets.

Here, we treat the strain design task as a network interdiction problem^22^ that maximally force cells to violate their wild-type phenotypes for production. That is to choose the optimum manipulations from *F* ^+^ ∪ *F* ^−^ ∪ *F* ^×^ in favour of biochemical production regardless of what the wild-type flux distribution is. As a result, we develop the following network interdiction problem:

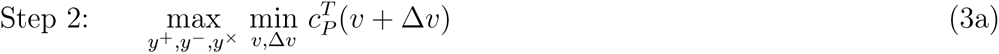

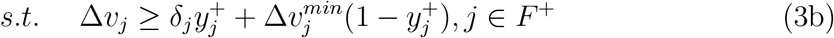

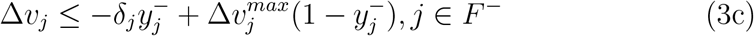

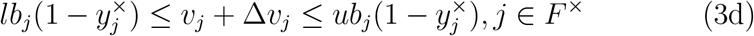

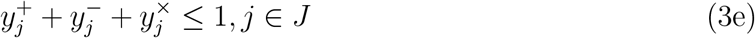

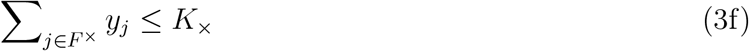

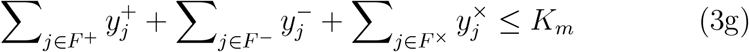

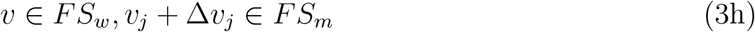

where *c*_*P*_ is a coefficient vector for the target biochemical. 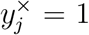 represents the knockout of reaction *j*, leading to zero flux in this reaction as illustrated by constraint (3d). Constraints (3f) and (3g) limits the allowable number of knockouts and the total number of manipulations.

The above statement is a special bilevel problem and can be formulated to a standard MILP using duality theory. ^9^ The resulting MILP can be handled either by a modern MILP solver or, if numerous alternative solutions are desired in a single run, by the hybrid Benders algorithm.^22^

The solution to problem (3) contains some regulation-associated binary variables that have a value of one, i.e., *y*_*j*_ = 1, for some *j* ∈ *F* ^+^ ∪ *F* ^−^. The integer values represent the reaction targets whose flux needs a change for biochemical production. In order to determine how they are to be regulated in experimental implementation, the following classification rule is applied:

reaction *j* ∈ Up-regulation Set ⟺ |*v*_*j*_| *<* |*v*_*j*_ + Δ*v*_*j*_| AND *y*_*j*_ = 1

reaction *j* ∈ Down-regulation Set ⟺ |*v*_*j*_| *>* |*v*_*j*_ + Δ*v*_*j*_| AND *y*_*j*_ = 1

The output of the model (3) predicts which reaction should be up- or down-regulated by at least the chosen flux change threshold. It does not impose exact fluxes on the mutant strain to guarantee the high production of target chemicals. In this sense, the resulting manipulations suggested by OptDesign could be experimentally more feasible than those obtained by existing tools.

### Computational implementation

OptDesign relies on model reduction and candidate selection for computational efficiency. Genome-scale metabolic (GEM) models can be significantly simplified by compressing linearly linked reactions and removing dead-end reactions (those carrying zero fluxes). Likewise, many reactions can be excluded from consideration with *a priori* knowledge that, for example, they are vital for cell growth or their knockout is not likely to improve target production. We followed the model reduction and candidate selection procedure^22,30^ (the detailed procedure can be found in the supplementary Figure 1), resulting in a candidate set of 342 knockout candidates for the latest *E. coli* GEM iML1515.^31^ The flux change threshold *δ*_*j*_ = 1 was used throughout the paper unless otherwise stated. **Algorithm 1** presents the pseudocode of OptDesign. It was implemented in MATLAB 2018b to be compatible with the Cobra Toolbox 3.0.^32^ All MILPs were solved by Gurobi 9.02.^33^ It is worth noting that a couple of minutes is enough for a modern optimisation software like Gurobi to identify a reasonably small set of up/down-regulation candidates. The source code is available for download at https://github.com/chang88ye/OptDesign.

#### Algorithm 1

The Overall OptDesign Procedure

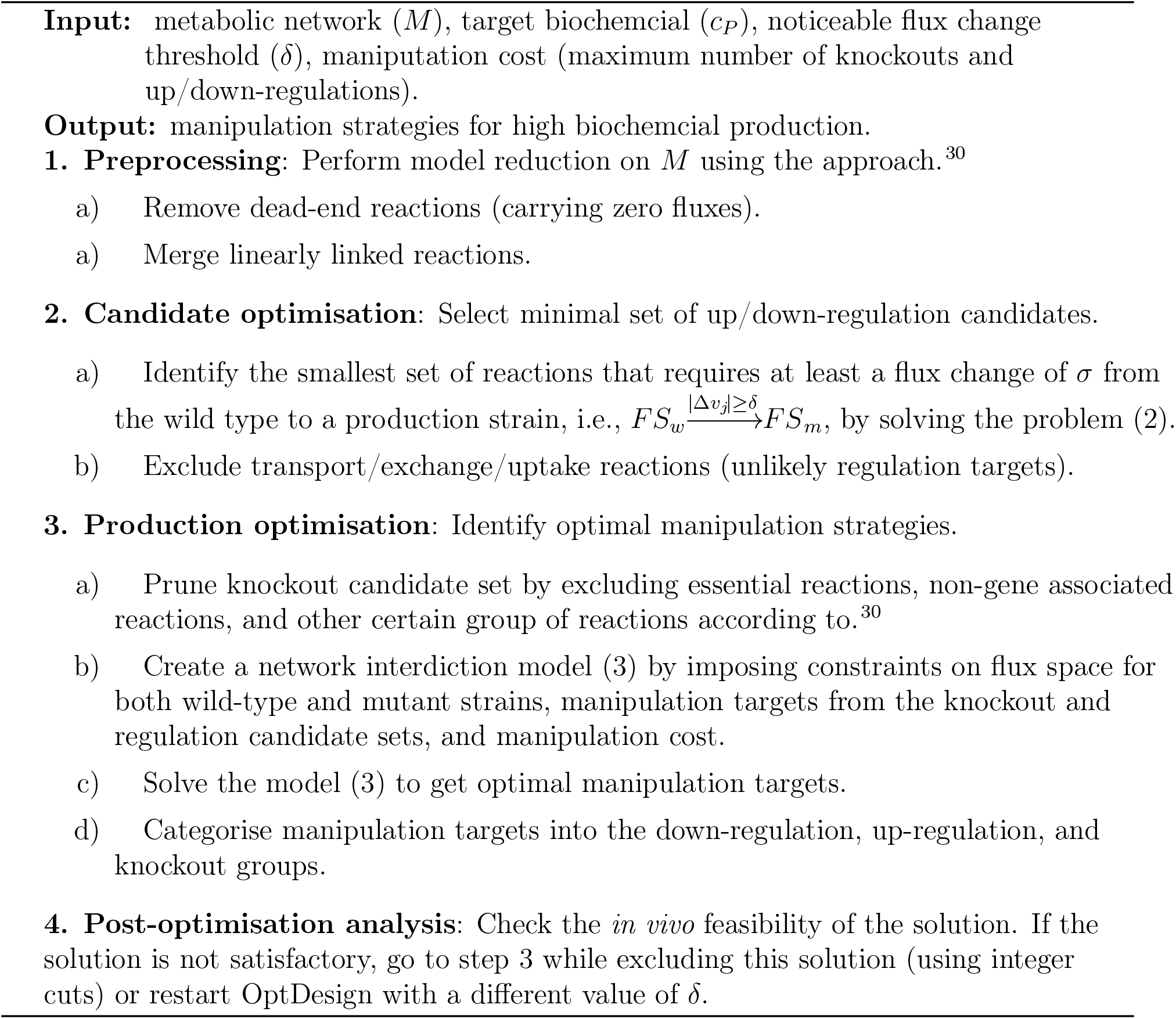

## Case Studies

The OptDesign framework was tested by identifying metabolic manipulations for the production of three industrially relevant biochemicals using the latest genome-scale metabolic model iML1515, ^31^ for *E. coli*. These biochemicals include the native succinate and non-native lycopene and naringenin, and their production has been extensively investigated through various strain design strategies.^26,34^ Similar to,^26^ succinate production was simulated under anaerobic conditions by setting the oxygen uptake rate to zero in iML1515. The heterologous biosynthesis pathway added to iML1415 for the production of two non-native biochemicals can be found in the supplementary Table 1 and Table 2. The growth conditions were the same as in iML1515 except a minimal cell growth of 0.1 *h*^−1^ was imposed on mutant strains for biochemical production. ^30^ At most 10 manipulations including no more than 5 knockouts were allowed, and the restriction on knockout is to intentionally favour gene expression manipulation over gene knockout. All optimisation problems in OptDesign were solved by Gurobi 9.02 on a MacBook with a 3.3 GHz Intel Core i5 processor and 16 GB RAM. The optimisation process was terminated by multiple stopping criteria whichever was met first, including time limit (10^4^ seconds) and optimality gap (5%). Indeed, we observed that the incumbent solution did not improve either after 3000 seconds or when the optimality gap reached 5%.

### Case study 1: Succinate overproduction

As a starting point, we wondered which reactions are likely to be good regulation targets and how they are distributed in metabolic networks. Therefore, we extracted the candidate regulation targets from the first step of OptDesign and reorganised them into different metabolic subsystems, as shown in Fig. 2. It can be observed that the majority of regulation candidates are from the Krebs cycle and fermentation products (i.e., formate, acetate and ethanol) that have the same precursor acetyl-CoA as succinate. It is suggested that all the reaction candidates from the Krebs cycle should increase their flux and those related to the formation of formate, acetate and ethanol should lower their activity. This prediction is consistent with many studies of succinate production.^26,35,36^ In additional to these two main subsystems, reactions from glucose metabolism, pyruvate metabolism and cofactor conversion/formation are also possible regulation targets. For example, the glucose transporter (GLCptspp) predicted for down-regulation here has been a deletion target in ^37^ to enhance succinate production.

**Fig. 2.**
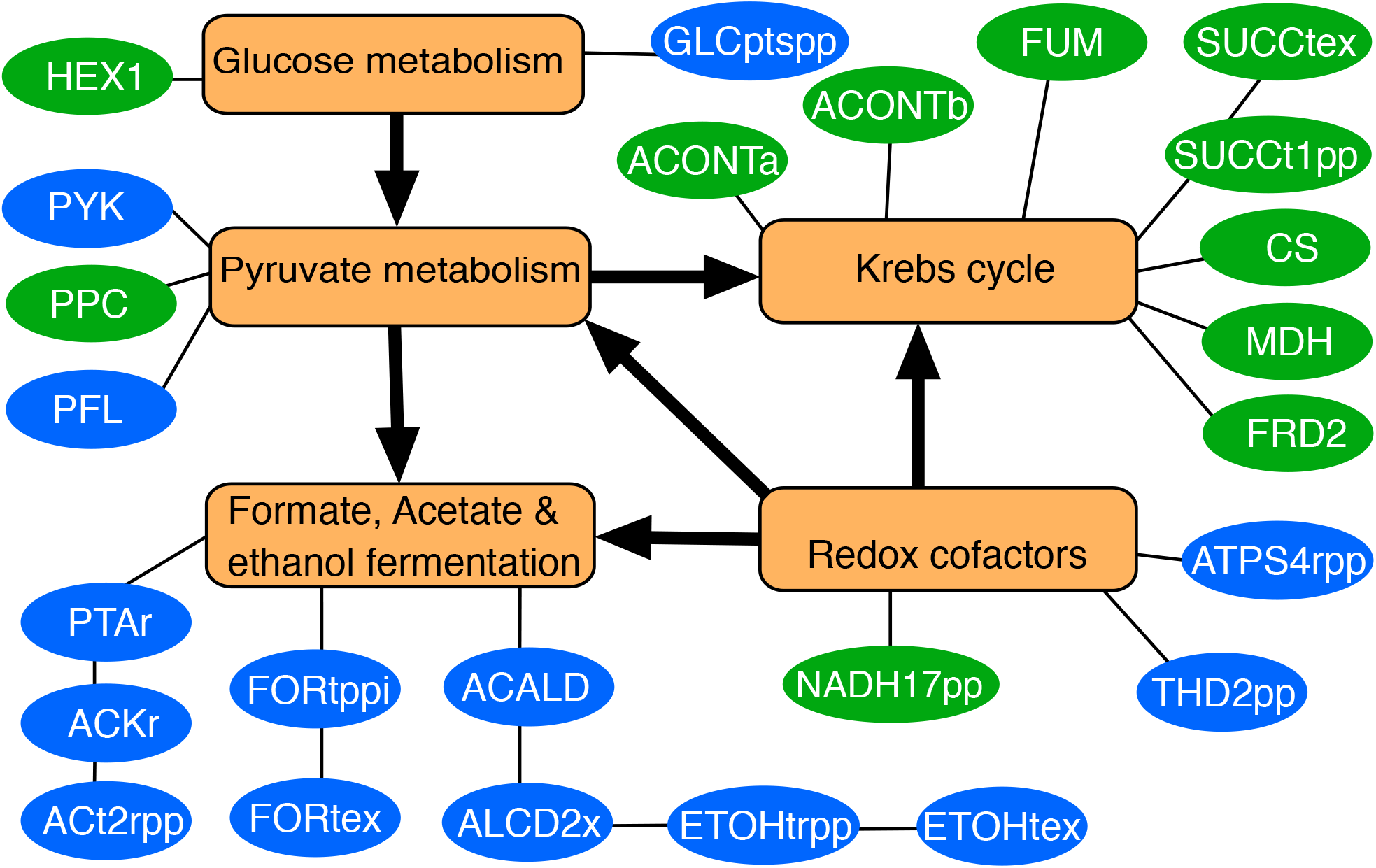
Reactions identified by OptDesign for up/down-regulation. Abbreviations of reaction names are borrowed from the iML1515 model definitions. Up-regulation and down-regulation reactions are in blue and green ovals, respectively. These reactions have been classified into different subsystems represented by orange rectangles.

Fig. 3a shows a few final design strategies identified by OptDesign that can improve succinate production. It suggests eight primary manipulations, including the knockout of five reactions, up-regulation of citrate synthase (CS) and pyruvate dehydrogenase (PDH), and down-regulation of periplasmic ATP synthase (ATPS4rpp). Two crucial enzymes in the formation of fermentation products lactate and ethanol, i.e., lactate dehydrogenase (LDH_D) and acetaldehyde dehydrogenase (ACALD), are suggested to be deactivated as they are considered as competing pathways consuming succinate precursors. These two manipulation targets have been observed in.^38,39^ The knockout of FAD reductase (FADRx) increases the availability of NADH, which has shown to be an effective approach to high succinate production.^40^ The methylglyoxal synthase (MGSA) pathway to lactate is another primary knockout target predicted by OptDesign. The removal of this minor pathway should result in pyruvate accumulation for succinate biosynthesis. Interestingly, this knockout has been implemented in,^36,39^ resulting in increased flux in the Krebs cycle. The ribulose-phosphate 3-epimerase (RPE) is another manipulation target identified by OptDesign, whose knockout forces D-xylulose 5-phosphate (xu5p-D) to be synthesised through the non-oxidative pentose phosphate pathway only. Therefore, it is expected that primary glycolytic flux flows into the precursors, e.g., phosphoenolpyruvate and pyruvate, of succinate.

**Fig. 3.**
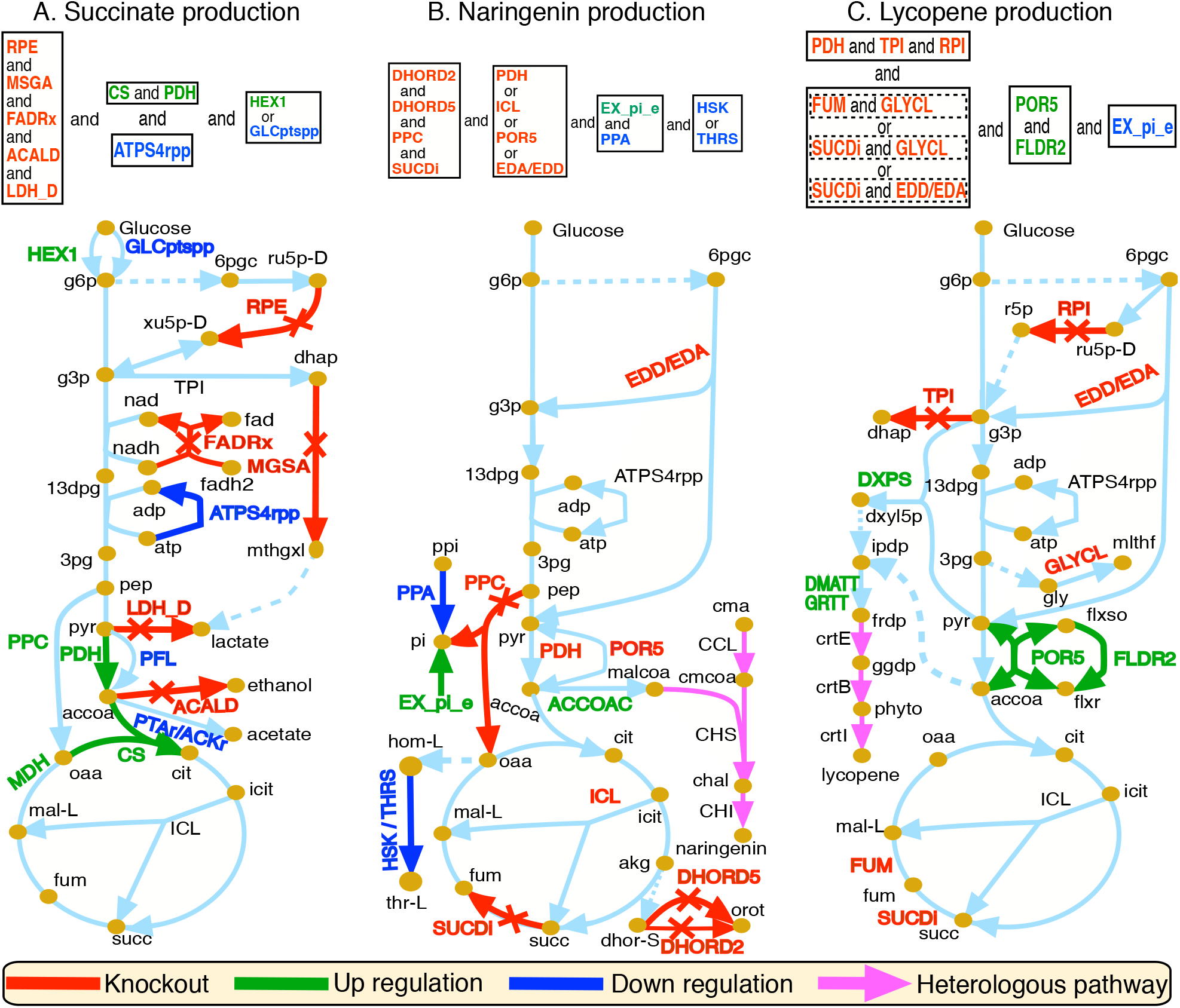
Design strategies identified by OptDesign for biochemical production in *E. coli*. Reaction names and their arrow symbols in same colour mean they must be manipulated in mutant strains. Reaction names coloured only (i.e., red, green, or blue) mean they are alternative manipulations. Dashed arrows represent a merge of multiple conversion steps to metabolites. Design strategies are summarised in boxes above the simplified metabolic maps. Abbreviations of metabolite names are as follows: g6p, glucose-6-phosphate; f6p, D-fructose 6-phosphate; g3p, glyceraldehyde-3-phosphate; 13dpg, 3-phospho-D-glyceroyl phosphate; 3gp, 3-phospho-D-glycerate; 6pgc, 6-phospho-D-gluconate; ru5p-D, D-ribulose 5-phosphate; r5p, alpha-D-ribose 5-phosphate; xu5p-D, D-xylulose 5-phosphate; dhap, dihydroxyacetone phosphate; mthgxl, methylglyoxal; pep, phosphoenolpyruvate; pyr, pyruvate; lac-D: D-lactate; dxyl5p, 1-deoxy-D-xylulose 5-phosphate; ipdp, isopentenyl diphosphate; frdp, farnesyl diphosphate; ggdp, geranylgeranyl diphosphate; phyto, all-trans-Phytoene; ppi, diphosphate; pi, phosphate; gly, glycine; mlthf, 5,10-methylenetetrahydrofolate; flxso, flavodoxin semi oxidized; flxr, flavodoxin reduced; accoa, acetyl-CoA; cit, citrate; icit, isocitrate; akg, 2-oxoglutarate; succ, succinate; fum, fumarate; mal-L, L-malate; oaa, oxaloacetate; hom-L, L-homoserine; thr-L, L-threonine; dhor-S, (S)-dihydroorotate; orot, orotate; malcoa, malonyl-CoA; cma, coumaric acid; cmcoa, coumaroyl-CoA; chal, naringenine chalcone; fad, flavin adenine dinucleotide oxidized; fadh2, flavin adenine dinucleotide reduced. Abbreviations of reaction names are referred1t5o the iML1515 model definitions.

OptDesign suggests to overexpress two enzymes in the succinate biosynthetic pathway, i.e., PDH and CS, which are intuitively straightforward to understand. In anaerobic *E. coli*, the PDH activity is either low or undetectable in order to maintain redox balance.^41^ However, it is observed that an *E. coli* mutant with activation of PDH for extra NADH improves succinate production.^39^ Overexpression of CS, which is also suggested in,^26^ has been observed to increase flux in the Krebs cycle in a malic acid production *E. coli* strain.^42^

OptDesign further predicts that high succinate production requires either up-regulation of glucokinase (HEX1) or down-regulation of a phosphoenolpyruvate-dependent phosphotransferase system (PTS) related reaction (GLCptspp), both of which have the same effect that glucose transport is favourably through the ATP-consuming HEX1 rather than the more efficient but phosphoenolpyruvate-dependent PTS route. Consequently, it improves the availability of PEP that is a precursor for biomass formation and many biochemicals including succinate. Both manipulation approaches have been observed to improve succinate yield.^35^ However, the increased ATP demand due to glucose transport via HEX1 has to be mediated by increased ATP production by other means. For this reason, OptDesign suggests down-regulation of ATPS4rpp to reduce cleavage of ATP to ADP in order to meet metabolic energy requirements. This prediction has been also suggested in.^43^

OptDesign also identifies a number of additional modification targets such as pyruvate formate lyase (PFL), phosphotransacetylase (PTAr) and acetate kinase (ACKr) that have been widely used as knockouts to increase flux towards the Kreb cycle in succinate studies.^36,39^ However, here OptDesign suggests to down regulate these enzymes instead of deactivating them completely. In addition, phosphoenolpyruvate carboxylase (PPC) and malate dehydrogenase (MDH) are also predicted as promising overexpression targets. This result is consistent with experimental studies that show increased succinate production through up-regulating these two enzymes. ^39,44^

### Case study 2: Naringenin production

A three-step pathway for naringenin was introduced into the metabolic network *E. coli* (see Fig. 3b) and unlimited coumaric acid (cma) was supplemented in the growth medium. OptDesign predicts that naringenin production requires four primary knockouts, one up-regulation and two down-regulations. The first two primary knockouts are dihydroorotic acid dehydrogenases (DHORD2 and DHORD5) that catalyse the oxidation of dihydroorotate to orotate in the pyrimidine biosynthesis pathway. The knockout of the underlying gene pyrD for these two reactions results in a reduced growth rate, ^45^ which might save carbon source for naringenin biosynthesis. The knockout of succinate dehydrogenase (SUCDi) creates a surplus of the biosynthetic precursor acetyl-CoA for naringenin, which has been experimentally observed in.^34^ Phosphoenolpyruvate carboxylase (PPC), a metabolic short-cut for the conversion of phosphoenolpyruvate to oxaloacetate and a byproduct phosphate, is also listed as a primary knockout. We postulate that, in addition to avoid the accumulation of phosphate, its deletion could not only direct flux through pyruvate to acetyl-CoA but also reduce the consumption of acetyl-CoA in the Krebs cycle for the mediation of oxaloacetate. In fact, PPC mutants were found to have a flux increase from pyruvate to acetyl-CoA in a ^13^C-labelling experiment.^46^ Two linear reactions, i.e., threonine synthase (THRS) and homoserine kinase (HSK), which are involved in the formation of L-threonine from L-homoserine, are predicted as down-regulation targets. This manipulation is expected to reduce carbon consumption in competing pathways, which therefore increases the carbon flux towards naringenin. Another down-regulation target is the inorganic diphosphatase (PPA) that catalyses the conversion of one ion of pyrophosphate to two phosphate ions. This manipulation is not intuitively straightforward and believed to create a combined effect with other manipulations to boost naringenin production. Since PPA down-regulation produces less phosphate which is needed in the added naringenin biosynthesis pathway, phosphate has to be balanced through an increase in its uptake channel, i.e., phosphate uptake reaction (EX_pi_e).

Aside from the above primary manipulations, it is also predicted that naringenin production strains must block at least one of the following reactions: two reactions on the Entner-Doudoroff pathway (EDD/EDA), pyruvate synthase (POR5), isocitrate lyase (ICL) and PDH. Blocking EDD/EDA might increase the use of glucolycosis, producing more ATP which is needed in the heterologous naringenin pathway. The removal of POR5 or PDH forces *E. coli* to use alternative conversion routes from pyruvate to acetyl-CoA without depleting coenzyme A (CoA), another primary precursor for naringenin biosynthesis. The knockout of ICL prevents the malate synthase reaction from consuming acetyl-CoA. In addition, OptDesign also predicts that the up-regulation of acetyl-CoA carboxylase (ACCOAC) helps to increase the production of naringenin, which has been implemented in. ^34^

### Case study 3: Lycopene production

A non-native lycopene biosynthetic pathway consisting of three key reactions were added to the metabolic network of *E. coli* (see Fig. 3c). A preliminary execution of OptDesign predicted the need of only one modification which is the overexpression of the gene encoding dimethylallyltranstransferase (DMATT) or the one encoding geranyltranstransferase (GRTT). While this manipulation intuitively makes sense, gene overexpression only in the upstream biosynthesis pathway of lycopene does not lead to high lycopene production due to low concentration of precursors, as experimentally illustrated in. ^47^ Therefore, we run our tool again while disallowing DMATT/GRTT to be valid regulation targets. Consequently, a variety of design strategies were identified, as shown in Fig. 3c. Specifically, all the design strategies are combinations of seven manipulations, consisting five knockouts, two up-regulations and one down-regulation. However, they differ from each other in only two knckout targets. The three primary knockouts, i.e., ribose-5-phosphate isomerase (RPI), triose-phosphate isomerase (TPI) and PDH, are linked to two precursors (i.e., glyceraldehyde-3-phosphate and pyruvate) of lycopene biosynthesis. The knockout of RPI reroutes the carbon flux flowing into the lycopene precursors using more effective metabolic routes (e.g., glycolysis) rather than the non-oxidative pentose phosphate pathway, which is consistent with the study of, ^3^ in addition to slowing down cell growth due to reduced ribose-5-phosphate formation for RNA and DNA synthesis. Both TPI and PDH knockouts should immediately increase the availability of the lycopene precursors, with the latter for increased lycopene biosynthesis being already confirmed experimentally in.^48^ Apart from glyceraldehyde-3-phosphate and pyruvate, acetyl-CoA is also an important precursor to form isopentenyl diphosphate, a building block for lycopene, using a different pathway. Therefore, it is expected that increasing the availability of acetyl-CoA should also improve lycopene. Unsurprisingly, POR5 is predicted as an up-regulation target in compensation for the loss of PDH for acetyl-CoA formation. Also, reducing the amount of acetyl-CoA flowing into the Krebs cycle was found to increase the flux towards isopentenyl diphosphate.^49^ This is fulfilled by either removing fumarase (FUM) or SUCDi in this study. Each of these two knockouts has to be paired with an additional knockout outside the Krebs cycle. This leads to three most frequent pairs, i.e., glycine leavage system (CLYCL) with FUM, CLYCL with SUCDi, and SUCDi with EDD/EDA. The predicted CLYCL knockout is believed to help reduce the cleavage of 3-phospho-D-glycerate into the glycine biosynthetic pathway so that more pyruvate can be accumulated. Alternatively, blocking the Entner-Doudoroff pathway allows more flux into glycolysis, leading to a higher production of the two precursors (i.e., glyceraldehyde-3-phosphate and pyruvate) for lycopene.

The NADPH-dependent flavodoxin reductase (FLDR2) is another primary up-regulation target predicted by OptDesign. Overexpressing FLDR2 is thought to balance the significantly increased ratio of NADPH to NADP^+^ caused by the last step of the lycopene biosynthetic pathway. Lastly, it is predicted that reducing the phosphate uptake rate improves lycopene production. This is probably because two out of the three reactions added for lycopene biosynthesis produce diphosphate that can be converted to phosphate, and a flux decrease in this uptake reaction rebalances phosphate in the system.

## Discussion

This paper has presented a new computational tool, called OptDesign, to aid strain development through rational identification of genetic manipulations including reaction knockout and flux up/down-regulation. This tool has been benchmarked via three case studies of different biochemicals, demonstrating its capability of identifying high-quality strain design strategies to improve biochemical production.

OptDesign predicts well in its first computational step a set of candidates that can be potentially used as experimental regulation targets, as shown in the succinate case. In a second computational step the algorithm further prunes this set to a realistically acceptable size while optimising biochemical production. Interestingly, many of the predicted manipulations have been experimentally implemented in previous studies. Taking succinate production as an example, ten out of fourteen manipulations (MSGA, ACALD, LDH_D, HEX1, PDH, PFL, PTAr/ACKr, GLCptspp, PPC, MDH) suggested by OptDesign have been employed in succinate-producing strains.^35,36,39,44^ Specifically, it has been shown that engineered *E. Coli* strains KJ060 and KJ073 produce succinate yields of 1.2–1.6 mol/mol glucose after removing completing pathways that lead to by-products ethanol, acetate, formate and lactate.^39^ These strains were developed through added acetate in culture media because the deletion of PFL causes acetate auxotrophy under anaerobic conditions.^36^ However, OptDesign suggests there is no need to completely deactivate PFL. Instead, down-regulating it avoids acetate auxotrophy while still achieving high succinate production. Additionally, it is observed that glucose transport favouring glucokinase over pep-dependent PTS yields higher succinate production.^50^ Furthermore, the overexpression of PPC in *E. coli* for increasing succinate yields has been confirmed in.^44^

OptDesign also suggests a few new modifications, such as the deletion of FADRx and RPE, the up-regulation of CS and down-regulation of ATPS4rpp, which to our best knowledge have not been experimentally implemented for succinate production. While up-regulation of CS has been shown to increase malic acid production, ^42^ it remains unclear whether this manipulation is also useful for succinate production. The suggested flux modifications on FADRx and ATPS4rpp reconfirm the importance of ATP and redox balance in succinate-producing strains.^39^ Deletion of RPE showed low flux in the Krebs cycle,^51^ suggesting that metabolic bottlenecks may exist upstream of the Krebs cycle. The design strategies predicted by OptDesign imply that a synergistic effect of RPE knockout with the other identified flux modifications can lead to a high production of succinate. Similar observations can be also found for the production of two non-native biochemicals naringenin and lycopene studied in this paper.

We have so far assumed that regulation targets can be selected only from the minimal regulation set derived from the first computational step of OptDesign. Under this assumption it ensures that regulation manipulations are used as few as possible, since suggested regulation levels cannot be exactly guaranteed in experimental implementation. However, in the case that multiple metabolic routes exist between two metabolites, the minimal regulation set will have only one of them included. In view of this, we have also computed the maximal regulation set by maximising the number of reactions that can have noticeable flux changes. Taking lycopene as an example, the number of regulation candidates increases sharply to 124 in the maximal regulation set from 47 in the minimal regulation set. Consequently, the resulting larger solution space makes it possible to identify design strategies with a better minimum guaranteed flux for naringenin (see Fig. 4).

**Fig. 4.**
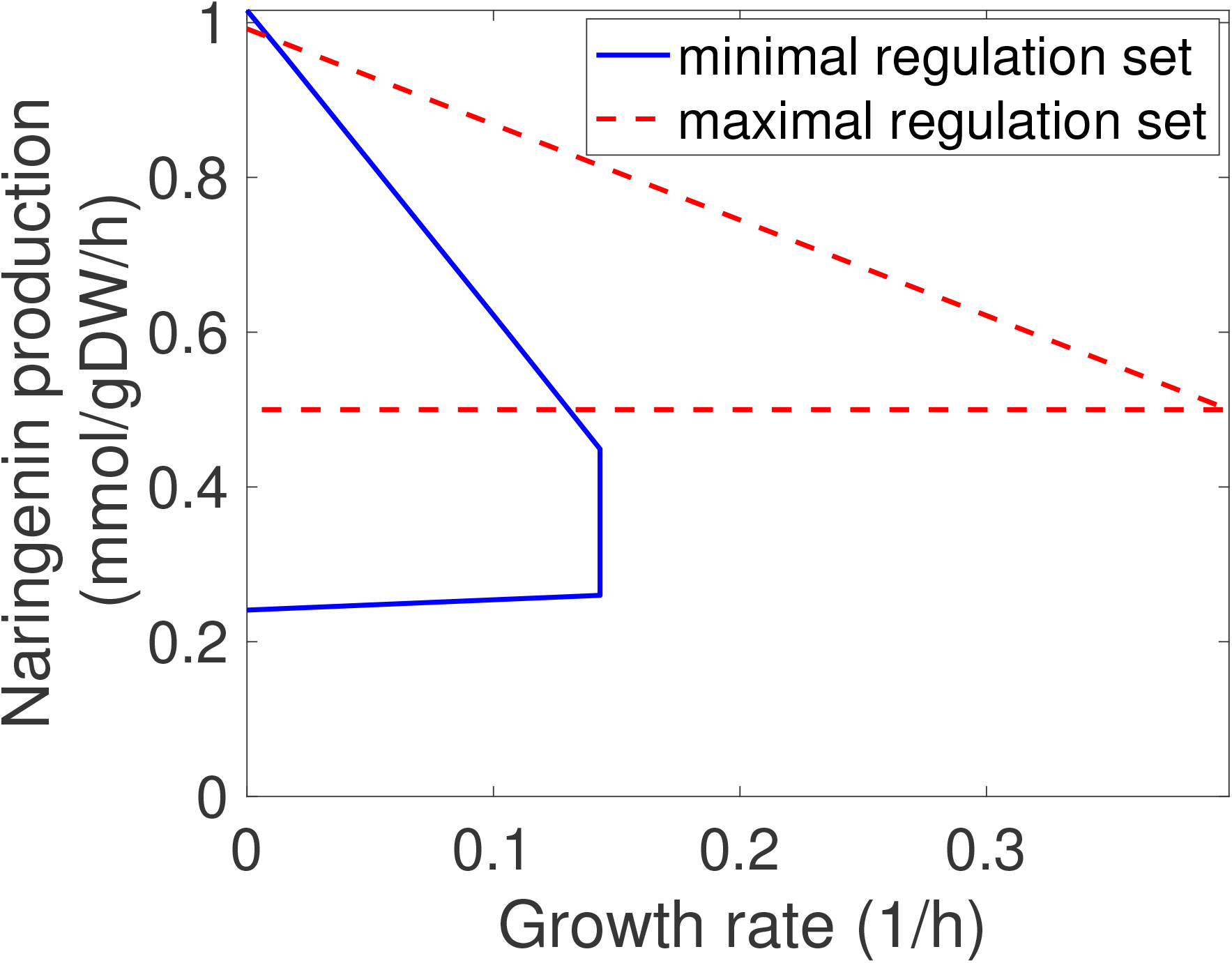
Production envelopes of different design strategies consisting of no more than 5 manipulations for naringenin. Production envelope illustrates the minimum and maximum production rate a production strain can achieve at different growth rates. The solid production envelope is for the design strategy using the minimal regulation set: FUM (knockout), ATPS4rpp (knockout) and DXPS (overexpressed), The dashed production envelope is for the design strategy using the maximal regulation set: FUM (knockout), IPDDI (overexpressed) and IPDPS (overexpressed). Reaction names is consistent with the genome-scale metabolic network model of *E. cli* iML1515.

In addition, the threshold *δ* defined for a noticeable flux difference between the wild-type and mutant strains influences the candidate regulation set, with a small threshold leading to a large number of regulation candidates. For example, the number of candidate reactions for regulation triples for succinate when the threshold drops to 0.1 from the current value 1.0 used in this paper (Fig. 5). The increased regulation candidates may present a higher chance of hosting optimal solutions for biochemical production but leads to a significantly larger network interdiction model that needs more computational costs to solve. Fig. 5 shows the influence of the threshold *δ* on succinate production. It is observed that the number of regulation regulations decreases as *δ* increases. However, the minimum guaranteed succinate production rate increases with *δ* until *δ* reaches 5, followed by a sharp decrease afterwards. The reason for the low production rate at *δ* = 10 is probably because some low-flux enzymatic steps whose regulation can significantly improve succinate biosynthesis are not included the small regulation candidate set. Thus, it may require careful selection of *δ* so that important regulation candidates are not excluded.

**Fig. 5.**
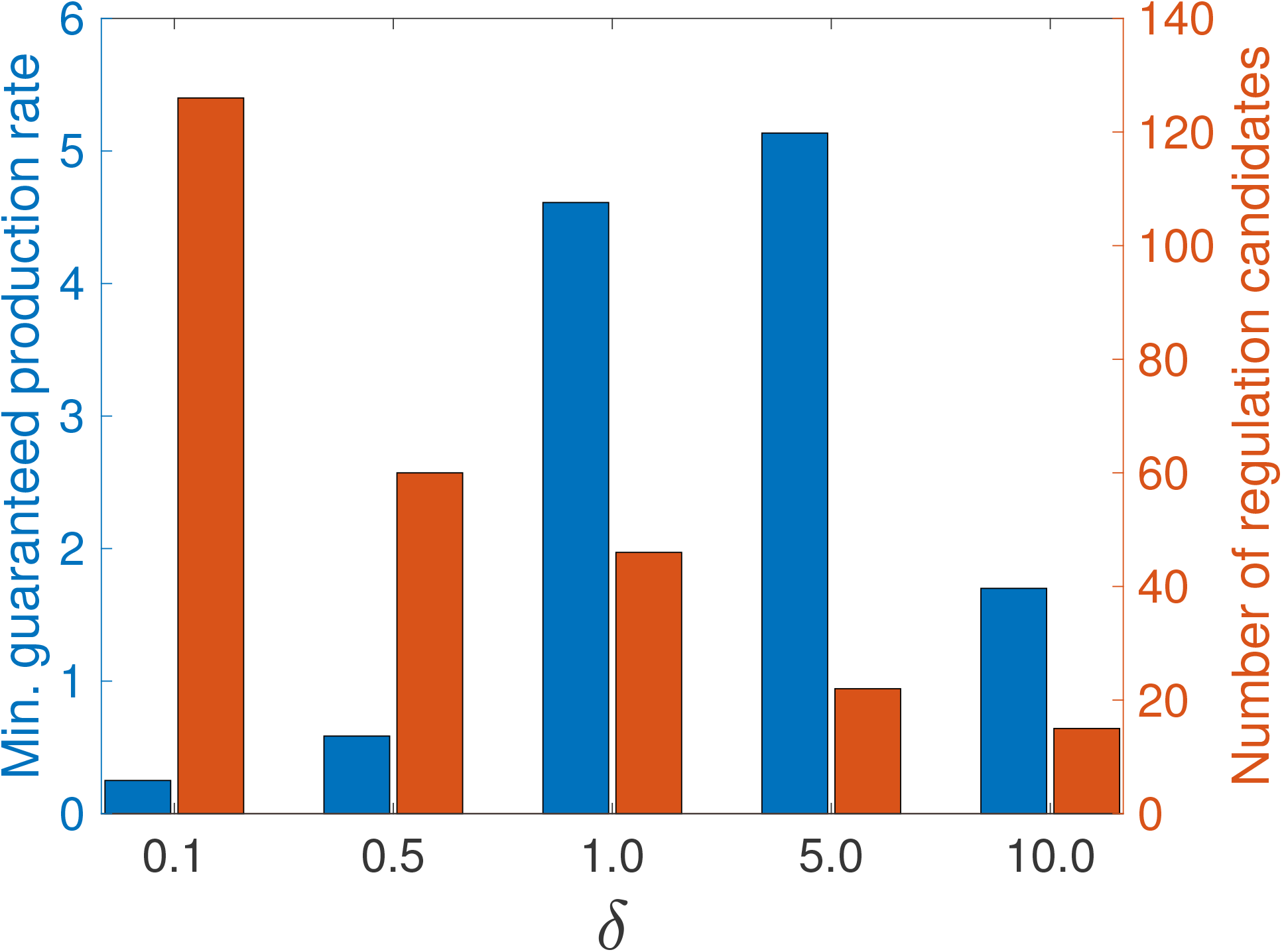
Influence of threshold *δ* on succinate production.

Furthermore, OptDesign highlights the benefit of flux regulation in strain design. For example, with the maximum number of 5 manipulations (including knockout and flux regulation) allowed, OptDesign found numerous design strategies for naringenin, with the best having a minimum guaranteed production flux of 1.73 mmol/gDW/h. In contrast, some existing strain design tools (e.g., OptKnock^9^ and NIHBA^22^) using knockout only did not identify any strategies leading to naringenin production. Like OptDesign, there also exist a few tools, e.g., OptForce^26^ and OptReg,^10^ that can identify both flux regulation and knock-out targets. We compared OptDesign with OptForce and OptReg in terms of manipulation targets. For succinate production (see Fig. 6), it is noticed that there is a large overlap between the design strategies predicted by these tools.

**Fig. 6.**
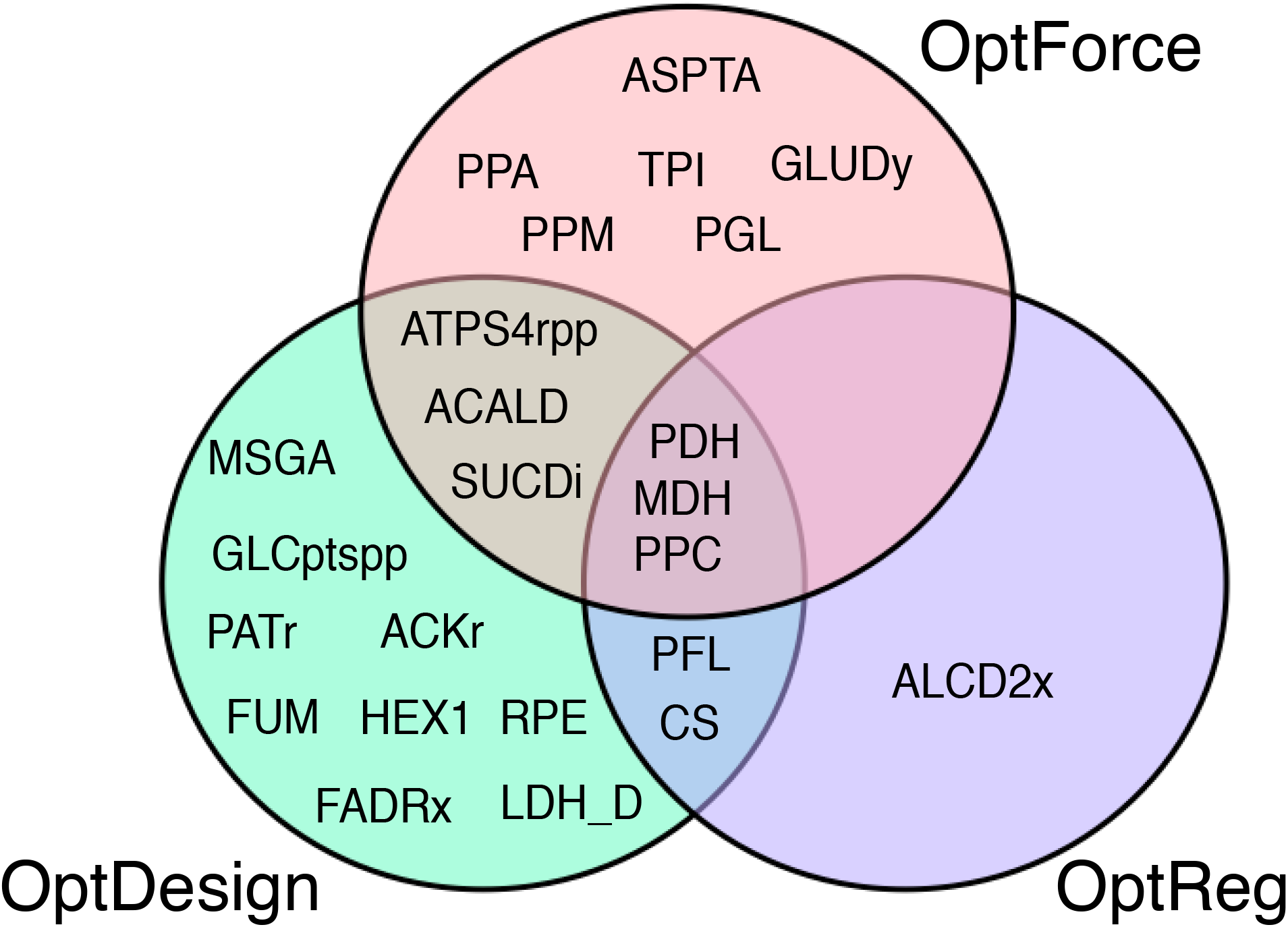
Comparison of intervention targets identified by different strain design tools. Reaction names is consistent with the genome-scale metabolic network model of *E. coli* iML1515. ments if available. Indeed, a measured wild-type flux vector can help refine the manipulation

Finally, OptDesign has been developed to identify metabolic manipulations regardless of whatever the wild-type flux distribution looks like, and it can be used with flux measure-candidates leading to a more accurate prediction of design strategies. In addition, the threshold for noticeable flux change defined in this work can be further adjusted with measured data, and different reactions can have distinct values for this parameter. Dedicated threshold values allow for a better prediction of rational flux modifications. Although OptDesign has been implemented for reaction-level phenotype prediction, it can be easily modified to predict design strategies at gene level. For example, OptDesign can be applied to metabolic network models with an advanced stoichiometric representation of gene-protein-reaction associations,^52^ from which design strategies consisting of gene targets can be identified.

## Acknowledgement

This work has been supported by the Engineering and Physical Sciences Research Council (EPSRC) for funding project “Synthetic Portabolomics: Leading the way at the crossroads of the Digital and the Bio Economies (EP/N031962/1)”. JRB acknowledges funding from MCIN/AEI/ 10.13039/501100011033 through grant PID2020-117271RB-C22 (BIODYNAMICS). NK is funded by a Royal Academy of Engineering Chair in Emerging Technology award.

